# Methodology, Validation and Testing of an Inexpensive Optical Planar-Motion Capture System

**DOI:** 10.1101/2025.07.18.665600

**Authors:** Joseph M. Mahoney, Jacob D. Weir, Courtney M. Jankowski, Benjamin W. Infantolino

**Author notes:** Corresponding author *Email address:* (Joseph M. Mahoney), *URL:* https://sites.google.com/view/jmmahoney/home (Joseph M. Mahoney). Alvernia University, Reading, PA 19601, USA.

## Abstract

Motion capture systems are ubiquitous in the gait analysis field. They are an excellent way to collect quantitative kinematic and kinetic motion data in an unobtrusive way. However, the cost for even a simple system is prohibitive to small research programs or clinics. Additionally, the required space, amount of equipment and set up time make them difficult to house permanently let alone move to different locations such as a clinic or the field. Here, we present a hardware and software solution to capture a more limited scope of kinematic data, specifically joint angles. The setup requires only one off-the-shelf camera and all data processing is done offline on a standard desktop or laptop. For this version, a treadmill was used to capture multiple strides. It required ten feet of clearance on the side for the camera and the hardware cost was approximately $300. The system is validated against an electrogoniometer and joint angles are found to be, on average, within 0.5^°^. The system is then employed in a small-scale case study of tracking knee and ankle angles for subjects walking with Ankle Foot Orthoses. We are able to detect clear differences in the ankle range of motion when comparing with-AFO to without. The knee and ankle joint profiles agree qualitatively with the literature.

## 1. Introduction

3D motion capture systems such as those from Vicon or Motion Analysis are the gold standard for capturing kinematic and kinetic data for gait anal-ysis. However, these systems have several disadvantages. A modest system for a research lab can cost $50,000 to $200,000. A minimum footprint of 5m *×* 5m is required for the system’s cameras and capture volume (Vicon, 2017) and the hardware is sensitive and difficult to transport. Finally, setting up and calibrating the system can take an hour or more. For a smaller research program, a classroom, or a clinic these issues can make a motion capture system impracticable.

For some applications, not all the information that these commercial systems collect is needed. In this article, we focus on the collection of joint angle data. One area where joint angles are pertinent is for Ankle-Foot Orthosis (AFO) fitting. Practitioners use their judgment and experience to determine the effectiveness of AFOs in achieving a more normal gait for affected patients. However, numerous studies have shown disagreement between practitioners when asked to describe the kinematics of patients with abnormal gait (Brunnekreef et al., 2005, Eastlack et al., 1991, Kawamura et al., 2007, Williams et al., 2009). An objective, qualitative assessment of abnormal ambulation would mitigate these discrepancies.

Alternative options for collecting motion data (including joint angles) exist but have drawbacks. Wearable devices such as electrogoniometers, accelerometers (Mayagoitia et al., 2002) and gyroscopes (Aminian et al., 2002) can encumber the wearer and only measure some joints well. Inertial sensors require initial positions and velocities to be known to calculate current position and the sensors drift over time.

Recently, low-cost systems have been developed to track joint angles using video cameras. Open source software was created to track markers in 3D (Figueroa et al., 2003) but requires multiple cameras and special markers. The Microsoft Kinect™ has been used as an off-the-shelf solution to motion capture (Fern’ndez-Baena et al., 2012, Gabel et al., 2012, Stone and Skubic, 2011). These methods are markerless but have not demonstrated the desired accuracy for joint angle measurement. A single-camera system was shown to track knee angle and gait events using markers (Yang et al., 2016). However, the accuracy of this system was shown to be within only 5^°^and tracks only one joint angle.

In this study, a simple and inexpensive hardware and software solution is introduced for tracking joint angles in a plane using off-the-shelf hardware and commonly-available software. The system is tested and validated for angle measures against an electrogoniometer. Finally, the system is tested with a small sample protocol to test its efficacy with subjects walking with AFOs.

## 2. Methods

This section describes the hardware setup and steps of the software to analyze joint angles. Here, lower limb analysis on a treadmill is used as the example to match the experimental protocol in Sec. 3. However, the system is not limited to lower limb analysis and could be employed for overground walking.

### 2.1. Data Collection and Analysis Methodology

To record lower limb motion, green pompoms (usually used for craft projects, hence referred to as “markers”) are affixed to the lateral side of the toe, heel, proximal and distal shank, and proximal and distal thigh of a subject. To increase the contrast between the background and markers in the video, a black sheet is placed behind subjects. The video is recorded 10 feet from the sagittal plane of the subject using a Canon r500 HD camera set to 1080p (1920 × 1080 pixels) resolution and 59.94 frames per second and saved as a mov file. A Matlab (R2016a, Mathworks, Natick MA) script was created to analyze the video after the data collection to determine the kneeand ankle angles.

First, the code opens the first frame of the movie and displays it to the analyst (see Fig. 1 (top)).

**Figure 1.**
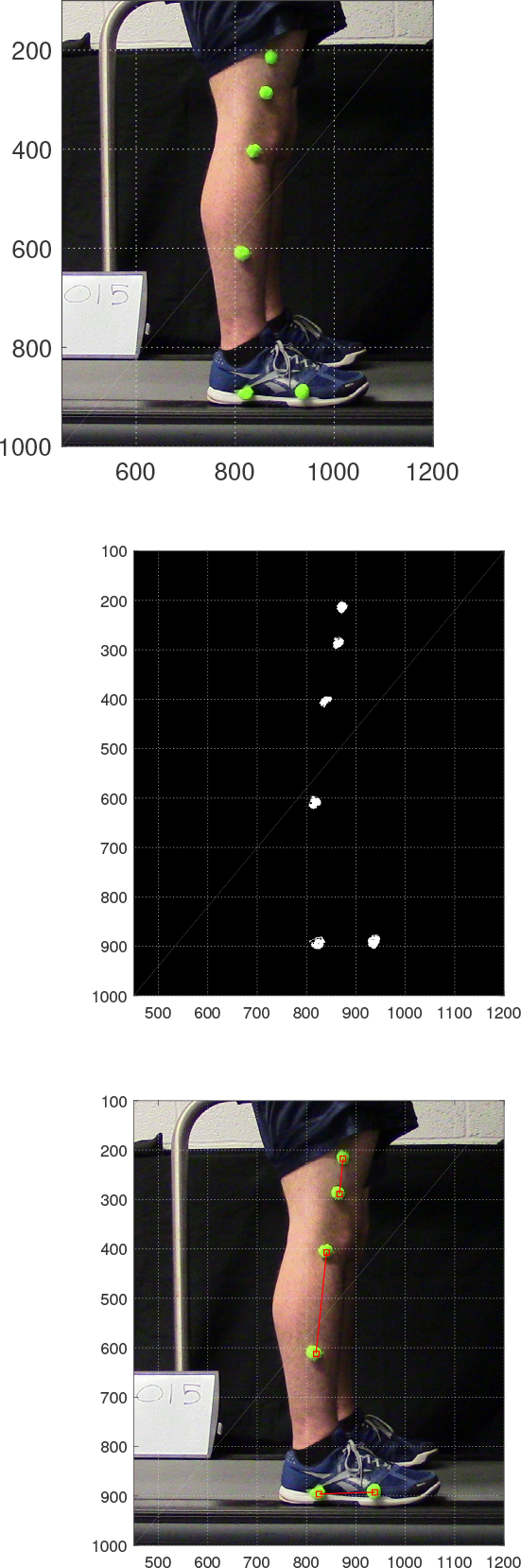
*Top*: Typical raw frame image taken from video. For clarity, it has been cropped to the area containing the legs and markers: it was originally 1920 *×* 1080 pixels. A black sheet was placed behind the subject to increase the contrast between the markers and the background. *Middle*: Image thresholded and then filtered to show only those pixels identified as markers (white). *Bottom*: annotated with the identified centroid locations of the markers (red squares) and connecting lines marking the limbs.

The marker locations are manually identified in the image by clicking on them with the mouse in proximal to distal order. The red-green-blue (RGB) values and *x*-*y* coordinates of the six clicked locations are stored. Due to differences in lighting and shadowing, the markers may appear to be slightly different colors in the video. The averages of the six red, green and blue values are taken to be the RGB color of a marker. Next, the frame image is thresholded. Here, the Euclidean distance between the RGB value for every pixel and the average RGB value of a marker is calculated. A binary matrix the same dimensions as the image is created. Every pixel in the image with a distance less than a set threshold has its corresponding location in the binary matrix set to 1 otherwise, it is set to 0 (see Fig. 1 (middle)). In other words, every pixel in the image is categorized as part of a marker or not. The value of the threshold can be adjusted based on the brightness of the video and the background contrast. If the value is set too small, pixels that are part of a marker will be ignored. If it the value is too high, parts of the background may be misidentified as being part of a marker. To reduce misidentified points and artifacts in the image, the binary image is filtered and small areas of pixels are removed. In Matlab, `bwareaopen` is employed. If a group of “1” pixels is below a set size threshold, they are then set to “0”.

This binary image is then put through a kmeans (Lloyd, 1982) clustering routine to locate the centroid of the six markers. Briefly, if it is known that there should be *k* distinct clusters of points in an image, the k-means process will select where to best place the *k* centroids. The `k-means` routine in Matlab was employed which allows additional options such as a starting guess as to the six centroid locations. The six manually-selected coordinates are used for this initial guess. The coordinates of the centroid of each marker are then identified. Figure 1 (bottom) shows a red square placed at the determined centroid location of each marker.

The process of finding the maker location on the subsequent frame now becomes automated. It is assumed that between frames the color of the markers varies little and the change in the position of a marker’s centroid is small. The RGB values of the next frame are loaded and then thresholded and filtered using the same rules as the first frame. The `k-means` routine is now run on this new image using the *previous* frame’s centroid locations as the initial guess for the new centroid locations. Using this initial guess reduces the processing time and increases the accuracy of the solution. This process repeats frame by frame for the entire trial.

In some frames, there may be shadowing or other artifacts that cause the routine to misidentify the proper centroid locations. The program checks if an identified centroid is too far from the centroids from the previous frame. When this is the case, the frame is dropped and is adjusted in the next step. This happens any time a marker is blocked in a frame, commonly occurring when the subject’s hand passes over the proximal thigh marker.

At this point, every frame has six centroid coordinates, but only in the first frame is each coordinate matched to the specific marker (e.g,. toe or distal thigh). Beginning with the second frame’s coordinates, the distances between a centroid and all the markers from the *previous* frame are found. Whichever marker has the smallest distance is taken to be the same marker in the current frame. This repeats for the other coordinates in the current frame and then the program moves to the next frame.

For the case of subjects walking on a treadmill, some identification shortcuts were taken. The topmost coordinate was always identified as the proximal thigh, the next coordinate down was always distal thigh, and the next down was always proximal shank. The vertical arrangement between distal shank, heel, and toe could change while walking so the previous method was used on only those points. This simplification reduced the total processing time, but for other motion, like running, this assumption may not be valid.

Using the coordinates of the markers, the 2D vectors of the rigid bodies (thigh, shank, and foot) are found by subtracting the distal point on the body from the proximal point. These vectors are shown as red lines on Fig. 1(bottom). The angles between the thigh and shank and between the shank and foot are then found using the dot product. On a body limb, the two markers are not necessarily affixed along the principal axis. To compensate for this, before or after the video is captured, a still calibration image is taken. In this image, the angle between the thigh and shank is held at 180^°^ (corresponding to 0^°^ flexion) and between the shank and foot is 90^°^ (corresponding to 0^°^ plantarflexion). The joint angles are calculated for this image and the values are stored as offsets for the calculated angles. The offsets are subtracted from the previously-calculated angles found from the recording, resulting in the true angle measures. Finally, the angle data is filtered by a 4^*th*^-order low-pass Butterworth filter with a cutoff frequency of 55 Hz to reduce high-frequency oscillations introduced by the marker identification process.

It should be noted that the threshold values for color difference, pixel group size, and maximum movement distance were selected based on the specific experiment. These values must be adjusted whenever the protocol changes. This adjustment takes the most time of the analyzer to ensure the best identification. Once set, the values were held constant for every subject and trial in the experiment for consistency. The program can handle a few dropped frames and fills in missing data points with a cubic spline fit (as done in commercial systems). However, having too many dropped frames will severely reduce the accuracy of the identification.

### 2.2. Goniometer Validation

To validate the optical system in Sec. 2.1, it was compared against measurements taken from an electrogoniometer (Delsys, Natick, MA). In this validation experiment, three markers were attached to the two ends and center on an analog goniometer. An electrogoniometer was then attached to the analog goniometer to monitor its angle. This device was filmed by the camera while the electrogoniometer simultaneously recorded. The goniometer was manually moved in ten-degree increments from 180^°^ to 130^°^ (the typical angle range for the knee) 10 feet from the camera. After closing and then opening once at this specified increments, the goniometer was then moved continuously between the limits.

The voltage output from electrogoniometer was calibrated to the known degree measure during the ten-degree increment phase by quadratic regression 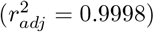. The video was calibrated by using the angle found in the first frame (180^°^) as the offset in every subsequent frame. The angle of the goniometer while in motion was calculated using the method in Sec. 2.1.

The electrogoniometer sampled at 5 kHz, so it was first down-sampled to match the camera’s sample rate of 59.94 Hz. The start times between the camera and electrogoniometer could not be synchronized electronically during the experiment. The goniometer was started before the camera and stopped after the camera. The two series were synchronized post hoc by finding the lag of maximum cross-correlation. This lag time was subtracted from the recorded time on the electrogoniometer to synchronize the devices. The angle measurements from the two devices while the goniometer is freely being opened and closed are shown in Fig. 2.

**Figure 2.**
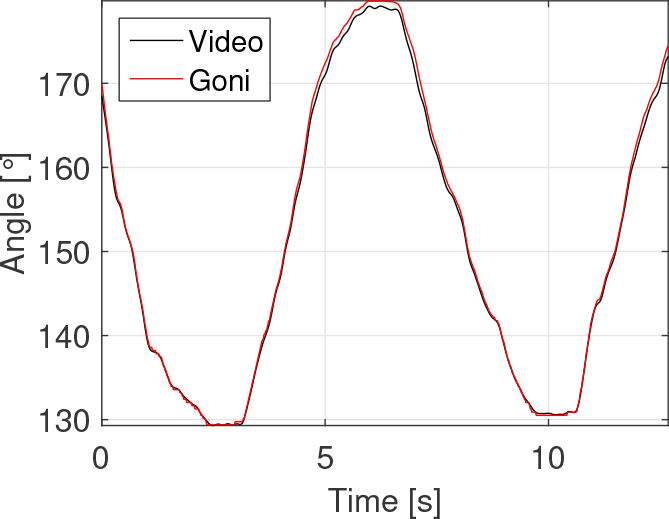
Comparison between “actual” angle (measured by electric goniometer) and measured angle (calculated from video). The times were synchronized by finding the lag of maximum cross-correlation. The mean difference between the two measures is *µ* = *−*0.453^°^ with the largest discrepancies happening at the extreme angle of 180^°^.

The difference in measured value between the goniometer and optical system was found to have mean of *µ* = *−*0.453^°^ [*−*1.77^°^, 0.404^°^] (mean [99% confidence interval]).

## 3. AFO Case Study

With the optical method shown to accurately (within 0.5^°^) measure planar angles in a controlled environment in Sec. 2.2, a small proof-of-concept experiment was run to test it with an experimental protocol on human subjects.

### 3.1. Methods

A convenience sample of four male and two female healthy subjects (*µ* = 22.12 years old, *σ* = 2.4) was recruited on the Pennsylvania State University Berks Campus. Any subject with a lower extremity injury or condition within the last 18 months was excluded. This study was done under the approval by the Penn State IRB.

Four walking conditions were used for the experiment: walking with (A) no AFO, (B) a spiral strut carbon fiber AFO (SAFO), (C) posterior strut carbon fiber AFO (PAFO), and (D) anterior strut carbon fiber AFO (AAFO). For each condition, subjects walked on the treadmill (Quinton Medtrack ST65, Quinton Instruments, Bothell, WA) at 1.2 m/s, a pace used in previous studies examining the effect of AFOs on normal gait kinematics and kinetics (Nair et al., 2010, Vistamehr et al., 2014), for 60 seconds to adjust to walking on the treadmill and with the AFO. Immediately following the adjustment period, the camera recorded walking for 90 seconds for each condition. The AFO was placed on the subjects’ right leg and the order of the four conditions was randomized for each subject. Shoes were worn during all conditions.

For each trial, the joint angles were calculated using the method outlined in Sec. 2.1. Subjects typically walked about 80 strides. Each stride was partitioned by taking data from one heel strike to the next. Because no force plate was used, heel strikes were determined by finding the times of maximum distance between the proximal thigh marker and the toe marker (Zeni et al., 2008). The time of each stride was normalized to 100% and resampled in 1% increments using linear regression. This made the joint angles a function of percent gait cycle rather than absolute time.

### 3.2. Results

For each subject and condition, the joint angles were averaged across percent gait cycle. A plot of a typical profile of joint angles for condition A (no AFO) is shown in Fig. 3. The solid line shows the mean angle and the dashed lines show one standard deviation above and below the mean.

**Figure 3.**
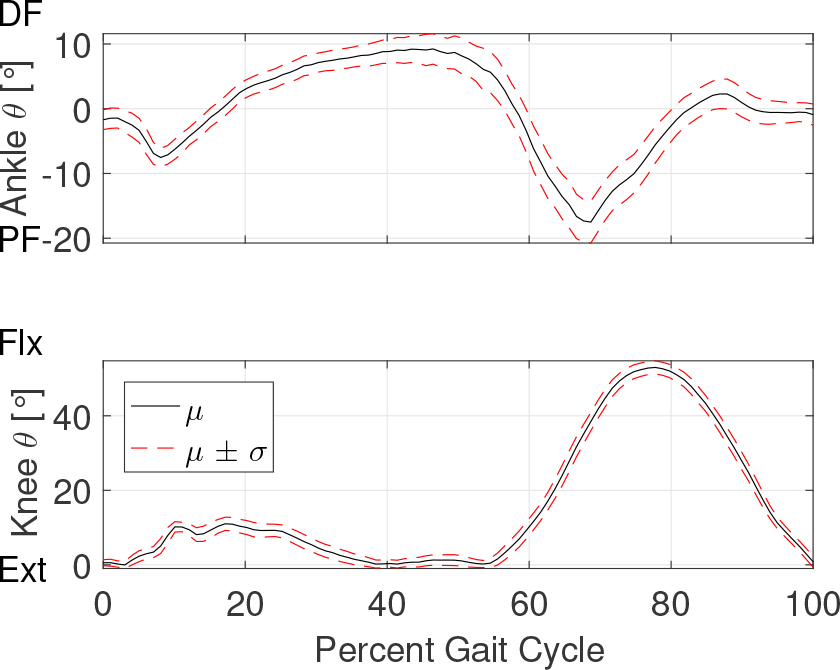
Typical ankle and knee angles over a cycle for one subject with no AFO. The black line indicates the mean joint angle and the dashed red lines are one standard deviation above and below the mean.

The angle profiles were then compared across walking conditions. A typical comparison of knee angles is shown in Fig. 4 (top) and ankle angles are compared in Fig. 4 (bottom).

**Figure 4.**
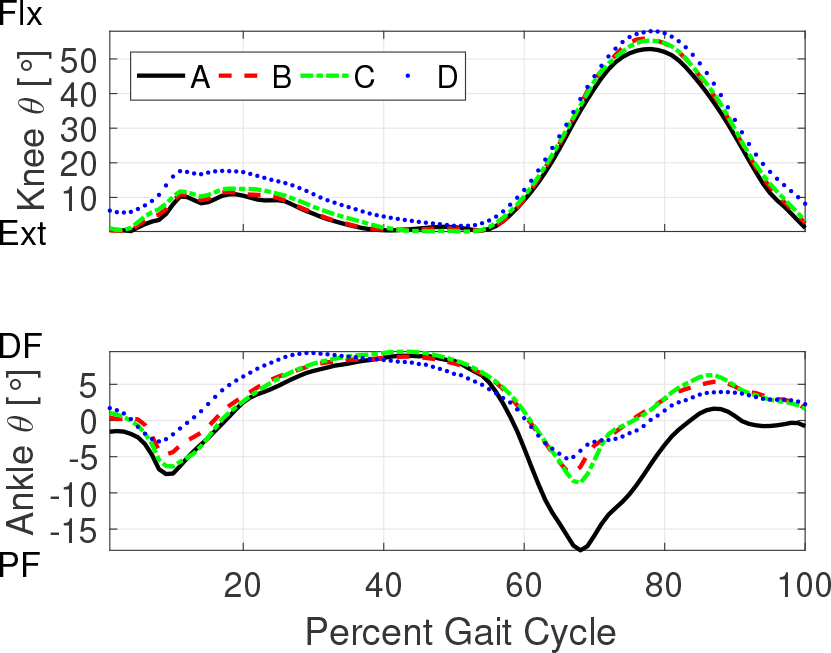
*Top*: Typical *mean* knee angle over a gait cycle averaged over all cycles for the four conditions for one subject. The vertical axis shows the amount of flexion (Flx) and extension (Ext) of the knee. Notice the maximum flexion in condition A is slightly smaller than the other three conditions. *Bottom*: Typical ankle angle over a gait cycle averaged over all cycles for the four conditions for one subject. The vertical axis shows the amount of dorsiflexion (DF) and plantar-flexion (PF). Notice the maximum plantar-flexion in condition A is much larger at 70% than the other three conditions with AFOs.

The range of motion (ROM) for the ankle and knee were calculated for each subject within each condition using the range of their mean angle profiles. A boxplot of the ROM values of the ankle across subjects is shown in Fig. 5 (top). The median without-AFO walking condition ROM was compared to each with-AFO condition’s ROM using a one-tailed matched sign test (Gibbons and Chakraborti, 2011). Because of the small sample size, it was not assumed that samples were normally, or even symmetrically, distributed around the median. Each analysis pair resulted in *p* = 0.0156. Thus, the median ROM for condition A was found to be significantly *larger* than the median for each with-AFO condition.

**Figure 5.**
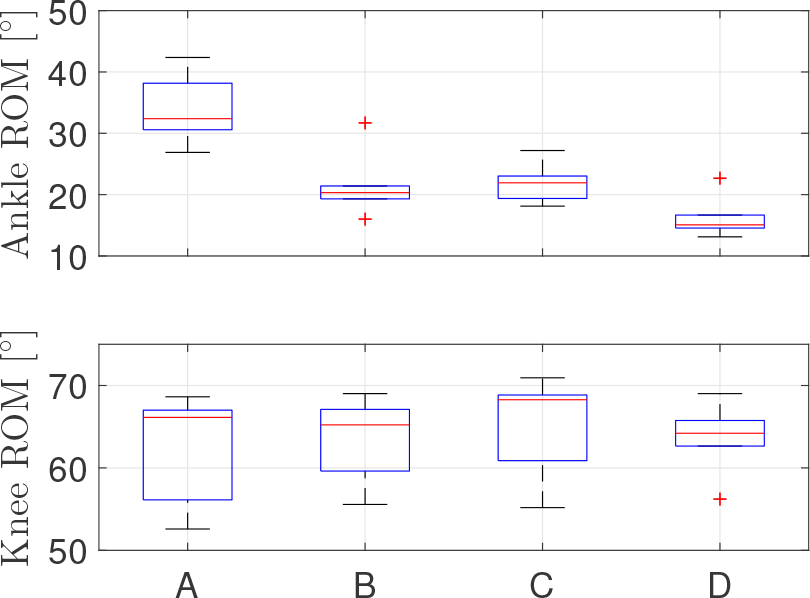
*Top*: Boxplot of range of motion of ankle. Whiskers show the range of the data within 1.5 IQR of the first or third quartile. Outliers are marked as asterisks but included in all analysis. Using a matched sign test, the median ROM of A is larger than the median of each other condition with *p* = 0.0156. *Bottom*: Boxplot of range of motion of knee. Using a matched sign test, the median ROM of A is not significantly different than each other condition with *p »* 0.05.

A boxplot of the ROM values of the knee across subjects is shown in Fig. 5 (bottom). The median without-AFO walking condition ROM was compared to each with-AFO condition’s ROM using a two-tailed matched sign test. Each analysis pair resulted in *p »* 0.05. Thus, the median ROM for condition A was not found to be significantly different than the median for each with-AFO condition.

## 4. Discussion

The validation test of the system against the electrogoniometer showed promising results. On average, the video calculated the angle only 0.5^°^ less than the “actual” value. Looking at the plot in Fig. 2 though, the largest discrepancies happened when transitioning from opening to closing the goniometer. The electrogoniometer itself may have been measuring inaccurately at this transition point. In future work, this system will be validated against a Motion Analysis system as a more reliable test.

The typical ankle and knee profiles for no AFO (Fig. 3) qualitatively looks like what is expected from the literature (Kadaba et al., 1990). There is variation between strides, which is to be expected, but the standard deviation looks reasonably small. If the system were inconsistently identifying markers, we would expect the deviations to be large.

The median ankle ROM was found to be significantly larger at 95% confidence for condition A than all the with-AFO conditions. This was expected as one purpose of the AFO is to restrict the ROM of the ankle while walking.

The median knee ROM was not found to be significantly different at 95% confidence between no-AFO and any with-AFO condition. However, per subject, differences in the motion of the knee are apparent (Fig. 5 (bottom)) to adjust for the restriction of the ankle joint.

## 5. Conclusions & Future Considerations

The optical system has been validated and produced angles comparable to measurements collected simultaneously with an electrogoniometer. The system was then employed in a test case with human subjects and delivered reasonable results. In followup work, this system will be compared against a Motion Analysis system for its validity on human subjects. The system is easily portable (a single camera and a sheet) compared to a full 3D system and does not occupy a large permanent footprint. The main expense for the system is the camera (approximately $300). Improvements such as 4K resolution and 120 fps recording will further increase the accuracy of this system. Compared to even the most budget-priced commercial 3D optical systems, this system is about 2% the cost.

A downside of this version is the time to process the files. On a standard desktop computer, the movie file took ten to fifteen minutes (wall time) per minute of video to calculate the joint angles. This may be sped up by parallel processing and a newer computer. In future versions, the software will be converted to an open source language, such as Python, to avoid costly software licensing and make the system more accessible. Changing to another language and optimizing the code will increase the processing speed.

In the case study presented in Sec. 3, only knee and ankle joints were measured. However, the software can be modified allow for more than six mark-ers to be used and therefore more than two joints and the upper body to be tracked. The system could also be used to capture a few strides of overground walking.

With the ease of portability, the system will be simple to move within rehabilitation centers to use with patients. As bilateral analysis is desirable in AFO analysis, the next iteration of this system will include two synchronized cameras that can be used to assess outcome measures such as symmetry. Pathological subjects will be recruited to walk with several AFOs. A clinician’s assessment of the best one will be compared to a quantified value from the joint angle analysis.

## Supporting information

Subject Demographics

## Acknowledgements

The authors would like to thank Virginia Gagliardi and Boas Surgical for use of their AFOs and helpful discussions. They would also like to thank the volunteer subjects for donating their time.

## Author Contributions

Programming - JMM

Experimental Design - JMM JDW CMJ BWI

Data Collection - JDW CMJ

Data Analysis - JMM JDW CMJ

Manuscript Preparation - JMM

Manuscript Editing - JMM JDW CMJ BWI

